# Existence and implications of population variance structure

**DOI:** 10.1101/439661

**Authors:** Shaila Musharoff, Danny Park, Andy Dahl, Joshua Galanter, Xuanyao Liu, Scott Huntsman, Celeste Eng, Esteban G. Burchard, Julien F. Ayroles, Noah Zaitlen

**Author notes:** These authors contributed equally to this work (Julien F. Ayroles, Noah Zaitlen). Present address: Encompass Bioscience Inc., San Francisco, CA, 94158, USA. Present address: Genentech, South San Francisco, 94080, USA. For correspondence (SM) or (NZ).

## Abstract

Identifying the genetic and environmental factors underlying phenotypic differences between populations is fundamental to multiple research communities. To date, studies have focused on the relationship between population and phenotypic mean. Here we consider the relationship between population and phenotypic variance, i.e., “population variance structure.” In addition to gene-gene and gene-environment interaction, we show that population variance structure is a direct consequence of natural selection. We develop the ancestry double generalized linear model (ADGLM), a statistical framework to jointly model population mean and variance effects. We apply ADGLM to several deeply phenotyped datasets and observe ancestry-variance associations with 12 of 44 tested traits in ~113K British individuals and 3 of 14 tested traits in ~3K Mexican, Puerto Rican, and African-American individuals. We show through extensive simulations that population variance structure can both bias and reduce the power of genetic association studies, even when principal components or linear mixed models are used. ADGLM corrects this bias and improves power relative to previous methods in both simulated and real datasets. Additionally, ADGLM identifies 17 novel genotype-variance associations across six phenotypes.

## Introduction

Many complex phenotypes differ dramatically in their distributions between populations due to genetic and environmental factors. Both broad^1,2^ and fine-scale^3^ population differences are central to epidemiology^4^, pharmacogenomics^5,6^, biomedicine^7^, and population genetics^8,9^. In the context of association studies, statistical correction methods for population structure, such as principal components^10^ and linear mixed models^11^, have helped identify thousands of loci associated with hundreds of complex traits^12^. This underscores the importance of understanding the causes and consequences of fine-scale population variation.

To date, studies of phenotypic differences between populations and statistical correction methods have primarily focused on variation in population means. As we demonstrate below, while studying fine-scale population structure in UK Biobank, we discovered that phenotypic variance, in addition to phenotypic mean, varies between populations. Such “population variance structure” (in analogy to “population mean structure”) can produce substantial phenotypic differences between populations and has major biological and statistical implications. For example, we recently showed for sex-biased diseases, even a small difference in a disease’s liability variance can double its prevalence between groups^13^. Various evolutionary models^14^ also suggest that changes in phenotypic variance allow populations to adapt quickly in response to environmental perturbations^15^.

Although the causes and consequences of phenotypic variance heterogeneity remain poorly understood, several factors could drive population variance structure. First, it can result from non-linear interactions among genotypes (i.e. epistasis). Admixture between genetically diverse populations can disrupt fine-tuned epistatic interactions, increasing phenotypic variance^16,17^. Similarly, gene-environment interactions^18^ (GxE) can induce changes in phenotypic variance when environmental exposures differ between populations. Secondly, population variance structure can emerge under additivity. Phenotypic variance itself is a genetically-controlled quantitative trait^19,20^, and as such the frequency of alleles associated with different levels of variability (vQTLs) may differ across populations. Here we also demonstrate, for the first time, that natural selection can directly induce phenotype-variance structure.

To identify and model population variance structure we develop the *Ancestry Double Generalized Linear Model* (ADGLM). ADGLM accommodates arbitrary phenotypic and covariate distributions while accounting for broad- and fine-scale population structure of phenotypic mean as well as variance. Recent work has shown that modeling ancestry-variance effects can reduce biases of GWAS test statistics^21,22^. However, these methods are limited to modeling binary responses^21^ or major population groups^22^. Other studies tested for genotypes associated with phenotypic variance (vQTLs)^23^, but did not model population-variance relationships^24–26^, which generates false-positives when population variance structure exists. We show via extensive simulations that ADGLM reliably detects phenotypic variance structure and is robust to several violations of model assumptions.

To examine the utility of our approach, we first test for population variance structure with ADGLM in several large human datasets. We discover ancestry-variance associations for 12 of 44 tested phenotypes in ~113K UK Biobank British-ancestry individuals and 3 of 14 tested phenotypes in ~3K Mexican, Puerto Rican, and African-American individuals. Additionally, we find 42 ancestry-variance associations in Mexicans of DNA methylation, an epigenetic mark associated with environment^27^, disease phenotypes^28^, and ethnicity^29^. We further illustrate the utility of ADGLM in the context of genetic association mapping and find that relative to linear regression with principal components, modeling population variance structure leads to an increase in power, both in simulated and real datasets. We release ADGLM as open-source R code.

## Material and Methods

### Phenotypic models

For a continuous phenotype *y*, we assume

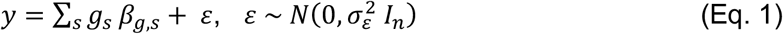

where *g*_*s*_ is the genotype of the *s*^*th*^ SNP, *β*_*g,s*_ is the genotype’s effect size, and errors in *ε* are assumed to be i.i.d. Gaussian. To model binary phenotypes, this model can be modified into a probit model by treating *y* as a liability and then thresholding. The main confounder in genetic association studies is population structure^30^. Linear regression with principal components (LR+PC)^10^ corrects for this by including the ancestry covariate *θ*:

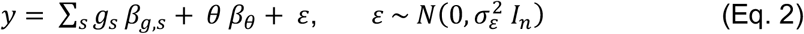

*θ* is often a matrix of genetic principal components, but it can contain ancestry admixture fractions or background covariates like age or sex. Linear mixed models (LMM)^31,32^ account for background genetic relatedness by including suitably normalized SNP genotypes, *Z*, as a random effect:

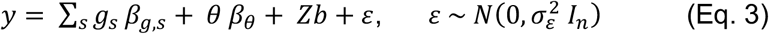

We ran LR+PC as ordinary linear regression for continuous traits and probit regression for binary traits. We ran LMM with pylmm^33^, choosing *Z* to have centered and scaled columns. Note that this LMM still models each sample as equally variable (modulo inbreeding).

### A statistical model for population variance structure

Population variance structure induces heteroskedasticity^34^, which violates standard linear model assumptions. Recent tests for heteroskedasticity^35,36^ or variance effects^24,25,37^ are appropriate when their assumptions are met, but they either cannot adjust for ancestry-variance effects, cannot simultaneously account for continuous ancestry and continuous phenotypic distributions^21,22^, or do not scale to UK Biobank sized cohorts. To jointly model phenotypic mean and variance, we develop a framework based on the double generalized linear model^38^ (DGLM), which has link functions and covariates for response mean as well as variance. Since we focus on ancestry-phenotype relationships, we call our framework the *Ancestry Double Generalized Linear Model* (ADGLM). The ADGLM uses standard estimates of ancestry (*θ*), such as fractional ancestry estimates or genetic principal components:

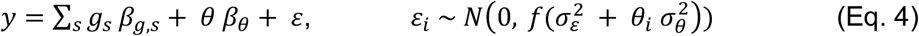

where *ε*_*i*_ is still entry-wise independent and *f* is the variance link function, typically the exponential function^39^. Negative variance effects decrease variance, but the exponential function guarantees that total variance is positive. The ADGLM accommodates dichotomous phenotypes via a probit link function or continuous phenotypes, as well as arbitrary phenotypic mean and variance covariates.

### Association testing with the ADGLM

The ADGLM framework enables likelihood ratio tests (LRTs) for ancestry and genetic associations. A 1 degree-of-freedom (df) test for ancestry-variance effect (i.e. population variance structure, 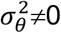) uses the null model *H*_0_ and alternative model *H*_2_, while a 1-df LRT for ancestry-mean effect (*β*_*θ*_≠0) uses models *H*_1_ and *H*_2_:

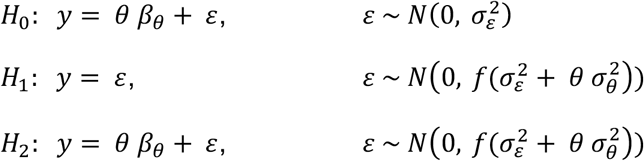

The ADGLM also enables 1-df association tests of mean genetic effect (*β*_g_≠0; *H*_2_ vs. *H*_3_) or variance genetic effect for vQTLs (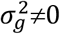; *H*_3_ vs. *H*_4_), both of which are corrected for population variance structure via *θ* and 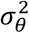. In addition, a 2-df test for mean genetic and ancestry-variance effects uses models *H*_0_ and *H*_3_:

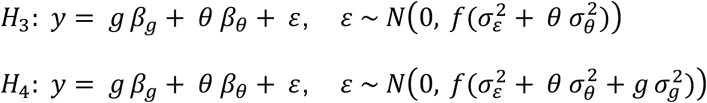

We used the R packages “dglm” and “glmx” for continuous and binary phenotypes, respectively, and the exponential variance link function throughout. Though we include a variance intercept term in continuous phenotypic models, we constrain it to one (1 = exp (0)) in binary phenotypic models to obtain identification. In the “dglm” package, standard errors of variance terms are approximated based on the leverages of the variance covariates^40^ and thus do not depend on the phenotype if it is scaled. These ancestry and genetic association tests, along with the diagnostic test for residual variance population structure, are implemented in the ADGLM code we released.

### Simulating data from a structured population

We simulated data from a structured sample of two population as follows. Ancestry of the *i*^*th*^ individual, *θ*_*i*_, is 1 for individuals from population 1, and 0 otherwise. We simulated the *s*^*th*^ SNP genotype of the *i*^*th*^ individual from population *j* as *g*_*is*_ ~ *Binom*(2, *p*_*sj*_, where *p*_*sj*_ is the SNP MAF in population *j*. We next simulated independent errors as 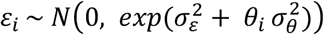 and phenotypes as *y*_*i*_ = *β*_*g*_*g*_*is*_ + *θ*_*i*_ *β*_*θ*_ + *ε*_*i*_. We took a sample of 200 individuals (100 per population), which has population variance structure when 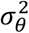 is non-zero. For the Figure 2 simulations with no genetic effect, m = 10,000 SNPs, *β*_*g*_ = 0, *β*_*θ*_ = 0, 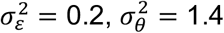. For the Table 1 simulations, m = 10,000 SNPs, *β*_*g*_ = 0.8, 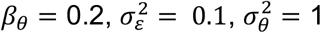. For the Figure 3 simulations with a true genetic effect, m = 10,000 SNPs, *β*_*g*_ = 0.6, *β*_*θ*_ = 0.3, 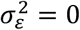 and 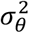 is one of 35 equally-spaced values between 0 and 2.

**Table 1:** Performance of genetic association tests applied to simulated data. We report false positive rate (∝ = 0.05, rows 1-7) or power (∝ = 5*e*^−7^, rows 8-12) of tests of genetic effect (*β*_*g*_ ≠ 0) for a range of MAFs (*p*_*1*_, *p*_*2*_) and genetic (*β*_*g*_), ancestry-mean (*β*_*θ*_), and ancestry-variance 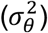 effects. The null hypothesis is true above the double line and false below it. Linear regression (LR) is miscalibrated in the presence of population structure. ADGLM is calibrated and powerful while linear regression with principal components (LR+PC) is often biased or underpowered.

| Row | Parameters |  |  |  |  | Discovery rate |  |  |
| --- | --- | --- | --- | --- | --- | --- | --- | --- |
| | $p_1$ | $p_2$ | $\beta_G$ | $\beta_\theta$ | $\sigma_\theta^2$ | LR | LR+PC | ADGLM |
| 1 | 0.5 | 0.5 | 0 | 0 | 0 | 0.051 | 0.051 | 0.054 |
| 2 | 0.05 | 0.5 | 0 | 0 | 0 | 0.054 | 0.052 | 0.054 |
| 3 | 0.5 | 0.05 | 0 | 0 | 0 | 0.054 | 0.050 | 0.050 |
| 4 | 0.05 | 0.5 | 0 | 0.2 | 0 | 0.140 | 0.053 | 0.054 |
| 5 | 0.5 | 0.5 | 0 | 0 | 1 | 0.051 | 0.051 | 0.055 |
| 6 | 0.5 | 0.05 | 0 | 0 | 1 | 0.074 | 0.086 | 0.053 |
| 7 | 0.05 | 0.5 | 0 | 0 | 1 | 0.032 | 0.021 | 0.057 |
| 8 | 0.05 | 0.5 | 0.8 | 0 | 0 | 0.993 | 0.783 | 0.725 |
| 9 | 0.5 | 0.5 | 0.8 | 0 | 1 | 0.698 | 0.693 | 0.848 |
| 10 | 0.5 | 0.5 | 0.8 | 0.2 | 1 | 0.702 | 0.702 | 0.850 |
| 11 | 0.05 | 0.5 | 0.8 | 0.2 | 1 | 0.467 | 0.205 | 0.606 |
| 12 | 0.5 | 0.05 | 0.8 | 0.2 | 1 | 0.838 | 0.269 | 0.135 |

### Simulating data from an admixed population

We simulated data from an admixed population composed of two source populations with a given *F*_*ST*_ as follows. We first drew the *s*^*th*^ SNP ancestral minor allele frequency as *p*_*s*_ ~ *U*(0.01, 0.5). We simulated source population MAFs with the Balding-Nichols model^41^ as *p*_*sj*_ ~ *Beta*(*p*_*s*_(1 − *Fst*)/*Fst*, (1 − *p*_*s*_)(1 − *Fst*)/*Fst*), *j* = 1, 2. The *i*^*th*^ individual’s population 1 ancestry fraction was drawn as *θ*_*i*_ ~ *U*(0.5, 0.9). For each allele (*k* = 1,2), we drew local ancestries as *γ*_*k*_~ *Bin*(1, *θ*_*i*_) and haploid genotypes as *l*_*k*_ ~ *Beta*(1, *p*_*sj*_), where *p*_*sj*_ is the MAF source population *γ*_*k*_. We formed the diploid genotype *g*_*is*_ of the *i*^*th*^ individual at SNP *s* as the sum of haploid genotypes. Finally, we simulated independent errors as 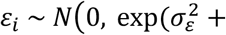 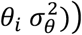 and phenotypes as *y*_*i*_ = *g*_*is*_ *β*_*g*_ + *θ*_*i*_ *β*_*θ*_ + *ε*_*i*_.

### Simulating data from an admixed population after differential selection

We simulated data as in “Simulating data from an admixed population” with the following modifications. For values of *T* between 0.05 and 0.25 in steps of 0.025, we simulated the *i*^*th*^ individual’s ancestry as *θ*_*i*_~ *N*(*T*, 0.15) and truncated it to (0, 1). We drew the *s*IJ SNP effect size as *β*_*gs*_ ~ *N*(0, 0.2). We then changed effect signs to induce a genetically-based correlation of phenotype and ancestry caused by three strengths of selection. Under neutrality, the sign of *β*_*gs*_ is unchanged, so *β*_*gs*_ is uncorrelated with ancestry. Under weak selection, the sign of *β*_*gs*_ is made positive with probability 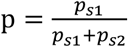 and negative with probability 1 − p, where *p*_*si*_ is ghijghk population *i* MAF. Since the Balding-Nichols model produces identical frequency spectra for all populations, p = 0.5, and *β*_*gs*_ and ancestry are perfectly correlated at half of the SNPs. Finally, under strong selection, the sign of *β*_*gs*_ is made positive if *p*_*s*1_> *p*_*S*2_ and negative otherwise, so *β*_*gs*_ and ancestry are perfectly correlated at all SNPs. These sign changes result in effect sizes 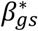. For the *i*^*th*^ individual, we simulated independent error as 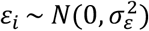 and phenotype as 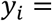 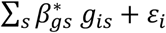. We did this for 2000 SNPs from a sample of 100 individuals for 1000 replicates.

### Outlier simulations

We simulated data from 1000 individuals for 1000 replicate simulations under the null 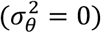 and alternative 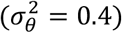 of the ancestry-variance test. For simulations with heavy-tailed errors, we simulated errors *ε* from the *t* distribution (df=6), simulated ancestry as *θ*_*i*_ ~ *N*(0.7, 0.4) truncated to (0, 1), and formed phenotypes as *y*_*i*_ = *θ*_*i*_ * 0.4 + *ε*_*i*_. In the simulations with real GALA II ancestry for *θ*_*i*_, we simulated errors as 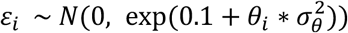) and phenotypes as *y*_*i*_ = *θ*_*i*_ * 0.4 + *ε*_*i*_. We transformed phenotypes or ancestry by inverse-variance quantile-normalizing them or truncating them to remove outliers more than two standard deviations from the mean.

### UK Biobank

We obtained UK Biobank data and restricted our analysis to ~113K British-ancestry individuals. We performed quality control steps as in a previous work^42^, resulting in genetic PCs and continuous phenotypes which are standardized to have mean 0 and standard deviation 1. We additionally quantile-normalized continuous phenotypes. For the variance association test, we adjusted for assessment center, genotype array, sex, age, and PCs 1-10 in the mean. We tested for variance effects (age, sex, PCs1-5) one at a time. The associated traits include ten blood traits, 15 disease traits, body mass index (BMI), blood pressure, educational attainment, basal metabolic rate, two measures of baseline lung function (forced expiratory volume in 1 second, FEV1, and forced vital capacity, FVC), age at menopause, hair pigment, skin pigment, and tanning.

### SAGE II and GALA II datasets

The Study of African Americans, Asthma, Genes & Environments (SAGE II)^43^ and Genes-Environment and Admixture in Latino Americans (GALA II)^44^ studies are comprised of admixed individuals (ages 8-21). Individuals were deeply phenotyped and genotyped. SAGE consists of 2,013 African Americans. The GALA II study consists of 4427 individuals, of whom 1245 are Mexican and 1785 are Puerto Rican. Genotyping resulted in 482,578 autosomal variants after filtering. We removed related individuals by excluding one of each of a pair of individuals with a REAP^45^ coefficient > 0.025, leaving 1160 Mexicans and 1612 Puerto Ricans. For both datasets, we removed genotypes with MAF < 0.05 and removed SNPs or individuals with more than 5% of genotypes missing. Global ancestry fractions were estimated with the program ADMIXTURE^46^ with two ancestral groups (Africans and Europeans) for SAGE and three ancestral groups (Native Americans, Africans, and Europeans) for GALA II.

### SAGE II and GALA II association testing

We tested the following phenotypes: asthma; allergy-related disease traits (eczema, hives, rhinitis, rash, and sinusitis); continuous traits (BMI, height); FEV1 and FVC; lung function changes after the first (*δ*_1_) and second (*δ*_2_) albuterol administrations. We also tested two skin pigmentation phenotypes: baseline melanin, the average of right and left body measurements of unexposed areas, and tannability, the difference between baseline and exposed melanin measurements^19^. For ancestry association tests, we included K-1 of K ancestry fractions in the variance model: for Mexicans and Puerto Ricans, we tested for African ancestry and included European ancestry as a variance covariate, and analogously for Native American and African, and African and European ancestry. For African-Americans (K=2), no additional ancestry variance covariate is required. For genetic association tests, we did not thin SNPs for LD nor impute missing phenotypic or covariate measurements. Where noted, genomic control^47^ was performed by dividing association test statistics by *λ*_*GC*_. We obtained GWAS associations of tested phenotypes from the NHGRI catalog^48^ on April 25, 2018 and thinned it to keep the strongest SNP association per locus, leaving 246 SNPs.

### GALA II methylation

We used QC for GALA II methylation data from whole blood as described in Galanter et al.^29^, resulting in batch- and cell type-adjusted methylation at 321,503 autosomal probes. Of the 124 Mexican individuals with methylation measurements, we removed those with outlier Native American ancestry (> 2 s.d. from the mean), leaving 117 individuals. We quantile-normalized methylation values and adjusted for age, sex, ancestry fraction, and asthma case status.

## Results

### Sources of population variance structure

Many studies have explored how genotype-by-environment interactions^18^ and epistasis^49–51^ may lead to a shift in phenotypic variance as a function of allele frequencies or environmental factors. Here, we consider another possibility: that differential selection between populations causes population variance structure under a purely additive model. To address this question, we simulated admixed populations that experienced differential selection.

We first generated allele frequencies at 2,000 SNPs from two ancestral populations under the Balding-Nichols model^41^. We then simulated effect sizes consistent with natural selection by correlating effect size and allele frequency difference between populations. We used a correlation of 0.0 under neutrality, 0.5 for weak selection, and 1.0 for strong selection. Finally, we simulated phenotypes using an additive model for a sample of 100 two-way admixed individuals composed of these ancestral populations with an average ancestry fraction, *θ*. Under neutrality, neither phenotypic mean nor variance depends on ancestry fraction (Figure 1). However, after either weak or strong selection, both phenotypic mean (Figure 1A) and variance (Figure 1B) depend on ancestry. This demonstrates that differential selection between populations is sufficient to induce population variance structure. In humans, strong, genetically-based ancestry-phenotype correlations are likely due to selection^52^, and may therefore be accompanied by population variance structure.

**Figure 1:**
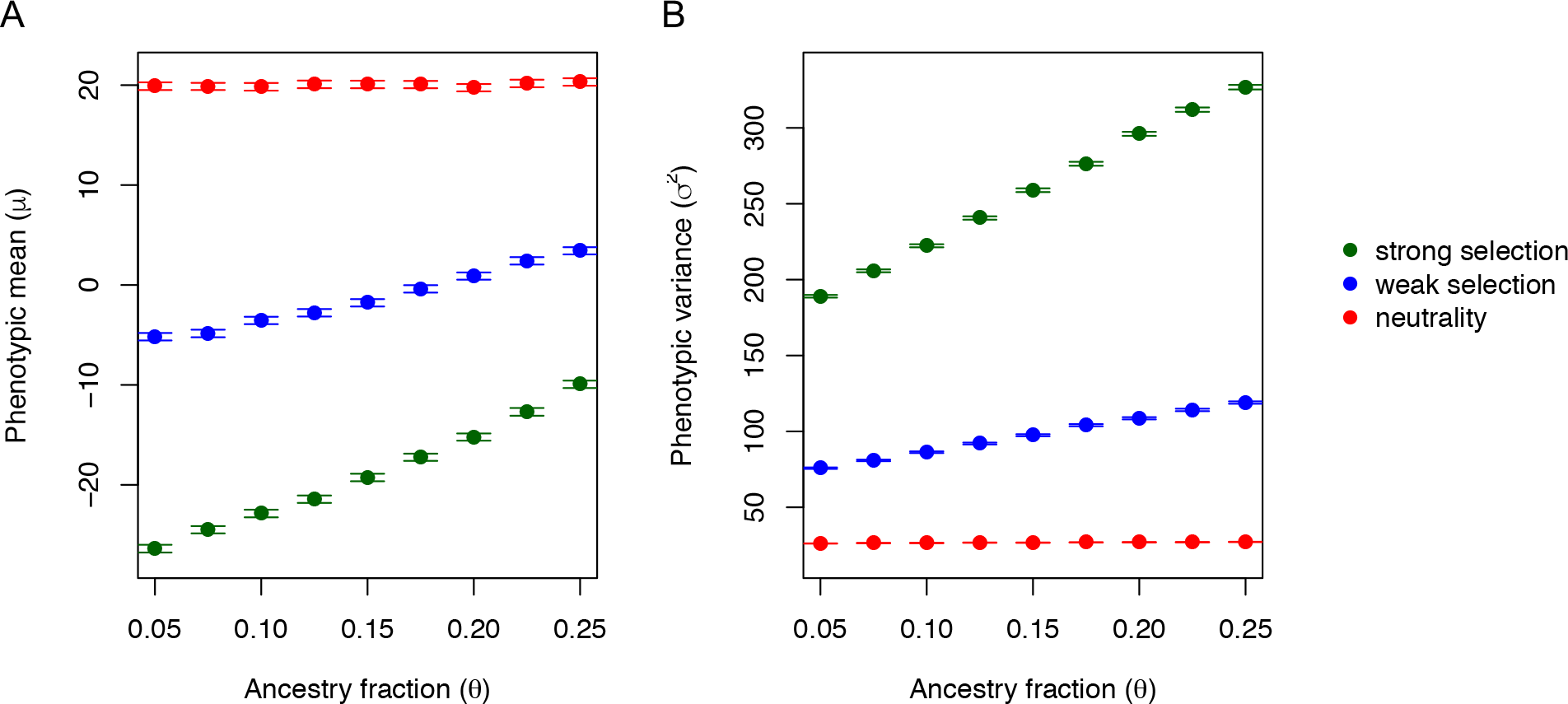
Differential selection induces population variance structure. (A) Phenotypic mean, *μ*, and phenotypic variance, *σ*^2^, of an admixed population with average ancestry fraction, *θ*, composed of two populations that experienced neutrality (red), weak selection (blue), or strong selection (green). Points are averages across 1000 simulations and bars denote 95% confidence intervals. Selection induces a dependence of phenotypic mean, as well as phenotypic variance, on ancestry.

### Ancestry-variance association tests

We first assessed the performance of the ancestry-variance test 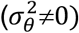 with ADGLM by applying it to simulated data from a structured sample of two populations (*P*_*1*_, *P*_*2*_) with MAFs *p*_1_ and *p*_2_. Since the MAF difference (*p*_1_ − *p*_2_) determines the genetic variance difference (2*p*_1_(1 − *p*_1_) − 2*p*_2_(1 − *p*_2_)) between populations, we considered three types of SNPs based on their MAF in the two populations: SNPs with a MAF difference that is large and negative (*p*_1_ = 0.05, *p*_2_ = 0.5), large and positive (*p*_1_ = 0.5, *p*_2_ = 0.05) and those with no MAF difference (*p*_1_ = 0.5, *p*_2_ = 0.5). We simulated 10,000 SNP genotypes and continuous phenotypes for 100 individuals from each population for a range of *β*_*g*_, *β*_*θ*_, 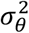 values. Under the null (no ancestry-variance effect 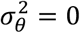), ADGLM is calibrated with a false positive rate of 0.052 when *β*_*θ*_ = 0 and 0.054 when *β*_*θ*_ = 0.2 at ∝ = 0.05 (also see Table S1). Under the alternative 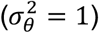, population *P*_*1*_ has greater phenotypic variance than *P*_*2*_, creating population variance structure in their combined sample. Here, ADGLM has power 0.463 when *β*_*θ*_ = 0 and 0.445 when *β*_*θ*_ = 0.2 at ∝ = 5*e*^−7^.

### Effect of population variance structure on genetic association tests

Genome-wide association tests commonly correct for population structure by using linear regression with principal components (LR+PC, Eq. 2) or linear mixed models (LMM^31^, Eq. 3 in Methods). We compared the performance of genetic association tests (*β*_*g*_≠0) with ADGLM, LR+PC, and LMM applied to data simulated as in “Ancestry-variance association tests”. First, we tested for a genetic effect on data simulated under the null (*β*_*g*_=0) with population variance structure, resulting in the quantile-quantile (Q-Q) plots in Figure 2. When population MAFs are equal, LR+PC is calibrated (Figure 2B, *λ*_*GC*_ = 1.01). However, when *P*_*1*_ MAF is greater than *P*_*2*_ MAF, LR+PC is inflated (Figure 2A, *λ*_*GC*_ = 1.41); when this MAF relationship is reversed, LR+PC is deflated (Figure 2C, *λ*_*GC*_ = 0.59). By contrast, ADGLM is calibrated for all MAFs: in Figs. 2A, 2B, and 2C, *λ*_*GC*_ is 0.98, 1.037, and 1.042, respectively. We also applied a standard LMM with ancestry as a fixed effect and the genetic relationship matrix as a random effect. LMM has the same miscalibration as LR+PC (Figure S1), so we do not consider it further.

**Figure 2:**
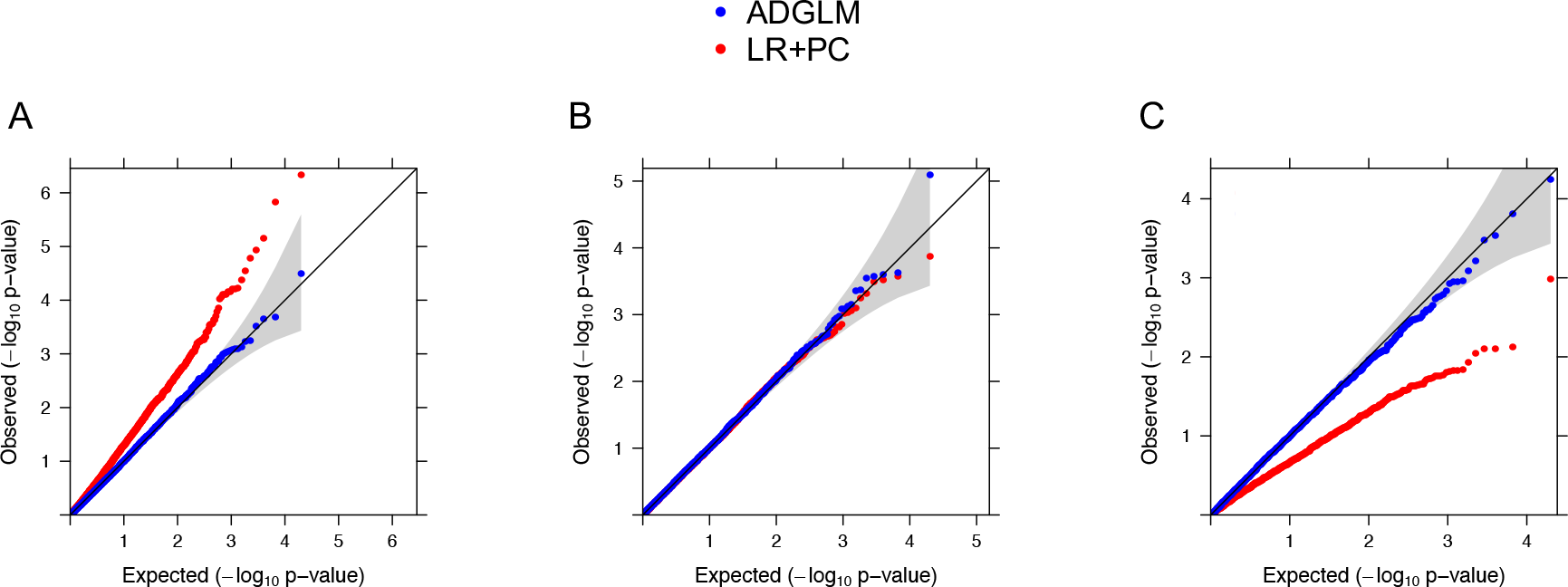
Q-Q plots of genetic association tests applied simulated null data. Data simulated with *β*_*g*_=0 and population variance structure for population MAFs: (A) *p*_1_ = 0.5, *p*_2_ = 0.05, (B) *p*_1_ = 0.5, *p*_2_ = 0.5, or (C) *p*_1_ = 0.05, *p*_2_ = 0.5. The null expectation is denoted by the black line and the null 95% confidence interval by the gray band. LR+PC −log_10_(p-values) in red are (A) inflated or (C) deflated when population MAFs differ; those from ADGLM in blue are calibrated.

Next, we assessed the performance of tests for *β*_*g*_≠0 on data simulated with a range of mean genetic (*β*_*g*_), mean ancestry (*β*_*θ*_), and ancestry variance 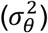 effects (Table 1, Table S1). When *β*_*g*_=0 and 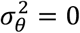, linear regression without ancestry adjustment (LR) is calibrated in the absence of population mean or variance structure (rows 1-3). However, LR is miscalibrated if there is population mean structure (row 4) or population variance structure and a MAF difference (rows 6-7). LR+PC and ADGLM perform similarly in the absence of population variance structure: they are calibrated (rows 1-4) and have similar power (row 8), despite ADGLM fitting an additional parameter. When there is population variance structure, ADGLM and LR+PC are calibrated if there is no MAF difference (row 5), whereas only ADGLM is calibrated if there is a MAF difference (rows 6-7). When *β*_*g*_≠0 and MAFs are the same, ADGLM is more powerful than LR+PC (rows 9-10). When MAFs differ, LR+PC has less power than ADGLM (row 11) or an elevated false positive rate (row 12).

Finally, we examined the power of genetic association tests (*β*_*g*_≠0) for varying ancestry-variance effects, 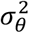. Power gains of ADGLM over LR+PC increase with 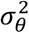 when MAFs are the same (Figure 3B) and when *P*_*1*_ MAF is less than *P*_*2*_ MAF (Figure 3C). When this MAF relationship is reversed, LR+PC has false positives, and ADGLM retains its power (Figure 3A). Taken together, these results demonstrate that tests for genetic association with LR+PC are miscalibrated and have false positives or false negatives when there is population variance structure while ADGLM is calibrated and powerful.

**Figure 3:**
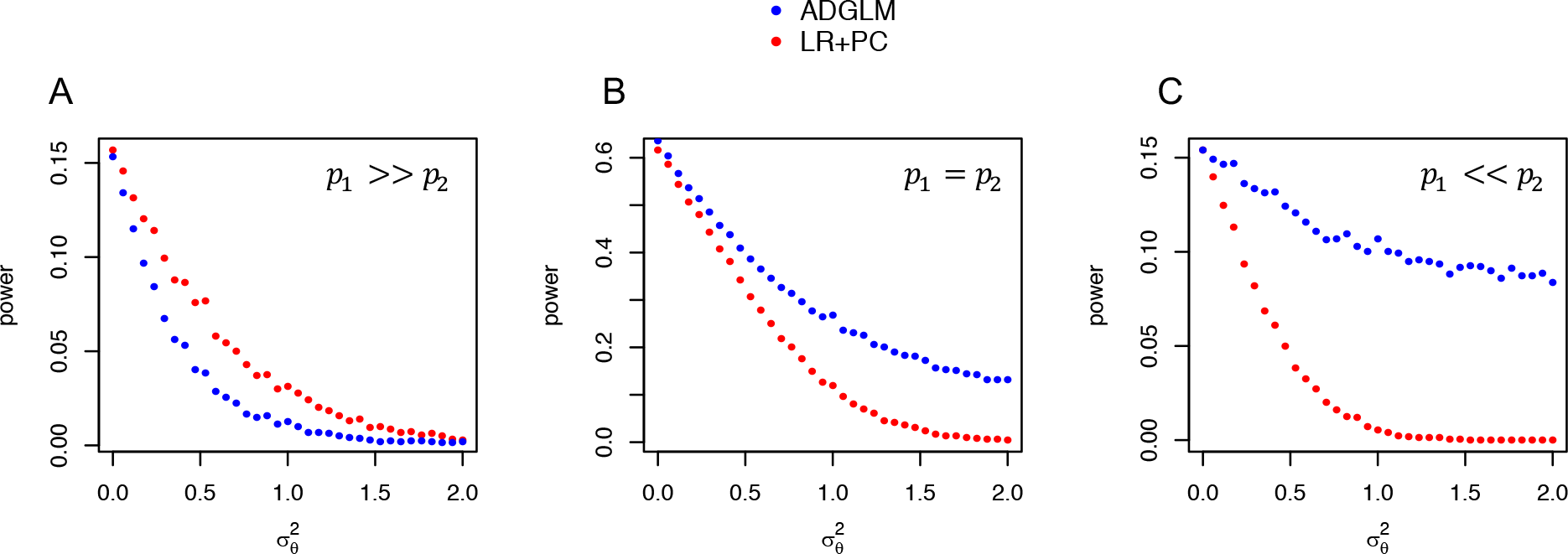
Power of genetic association tests applied to simulated data. Power (∝ = 5*e*^−8^) of tests applied to data simulated with varying 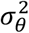 for three MAF cases. (A) For *p* = 0.5, *p* = 0.05, LR+PC is enriched for false positives, and ADGLM is well-powered. For (B) *p*_1_ = 0.5, *p*_2_ = 0.5 and (C) *p*_1_ = 0.05, *p*_2_ = 0.5, ADGLM has more power than LR+PC when there is population variance structure 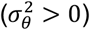.

### Diagnostic test for population variance structure

As we showed above, GWAS performed with standard corrections for population structure may result in biased test statistics in the presence of population variance structure. We developed a test for this bias that can be applied to GWAS summary statistics (Supp. Materials). It regresses association test statistics on the difference of expected genetic variances and tests for a non-zero slope, which occurs when there is residual population variance structure. This diagnostic test is well-powered on test statistics from LR+PC applied to data simulated with population variance structure (*p*=3.1×10^−26^, Figure S2) and is implemented in the ADGLM code repository.

### Sensitivity of ancestry-variance test to model assumptions

Double generalized linear models, like most linear models, assume regression errors are normally distributed^23^. We assessed the robustness of testing for 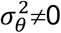 with ADGLM to violations of this assumption. We examined the ability of two transformations to reduce Type 1 errors under model misspecification: inverse-quantile normalization (“normalization”) and outlier removal (“truncation”).

We simulated data under the null with heavy-tailed errors (*t*-distribution, df=6) and applied ADGLM. Although ADGLM is miscalibrated (λ_GC_=1.22, FPR=0.079), phenotype truncation (λ_GC_=0.95, FPR=0.049) or normalization (λ_GC_=0.97, FPR=0.048) recovers calibration (Figure 4A). We next applied ADGLM to simulated data where 90% of replicates are null and found that relative to the original data (TPR=0.344, FPR=0.774), truncation (TPR=0.291, FPR=0.049) or normalization (TPR=0.315, FPR=0.048) improve both power and false positive rate.

**Figure 4:**
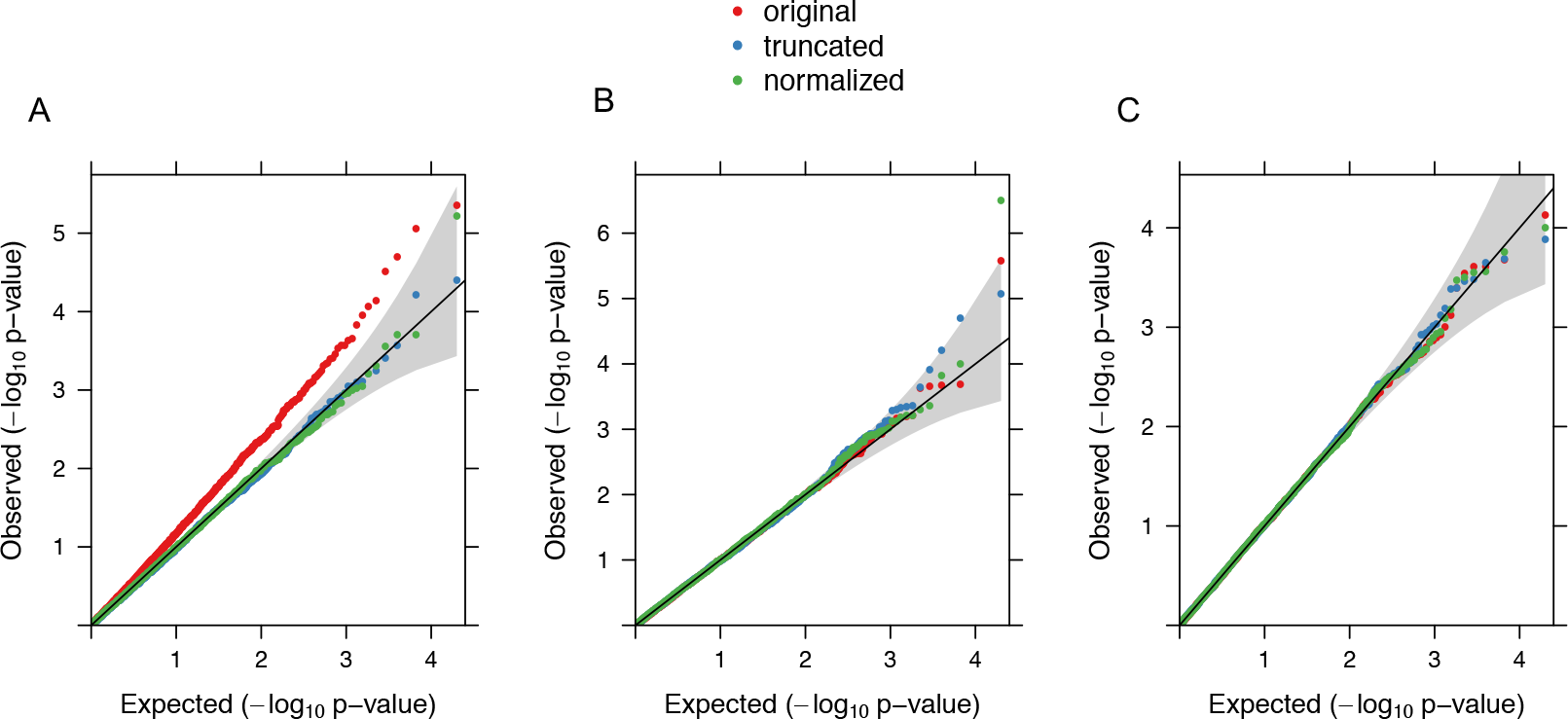
Effect of phenotype and ancestry outliers on ancestry-variance tests. Q-Q plots of ancestry-variance association tests applied to data simulated under the null 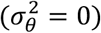. (A) Phenotype transformation reduces false positives of data simulated with heavy-tail, t-distributed error. (b, c) Tests applied to data simulated with real (B) African ancestry from Puerto Ricans or (C) European ancestry from Mexicans are calibrated and unaffected by ancestry transformation.

Next, we assessed the robustness of tests for ancestry variance effect 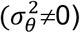 to non-normal ancestry distributions. We simulated phenotypes using ancestry fraction found in the Gene-Environment studies of Asthma in Hispanic/Latino children (GALA II^43^, Figure S3) as described in Methods. We first used the skewed African ancestry distribution from Puerto Ricans, where 2.0% (35) individuals are ancestry outliers. Applied to data simulated under the null, ADGLM is calibrated (λ_GC_=0.991, FPR=0.047) and minimally affected by ancestry truncation (λ_GC_=1.013, FPR=0.048) or normalization (λ_GC_=1.025, FPR=0.051) in Figure 4B. On data simulated under a mix of the null and alternative, performance is similar for original (TPR=0.067, FPR=0.047), truncated (TPR=0.057, FPR=0.047), and normalized data (TPR=0.076, FPR=0.050). Applied to data simulated with the bell-curved European ancestry from Mexicans with only three ancestry outliers, ADGLM is calibrated under the null (λ_GC_=1.00, FPR=0.050) and minimally affected by ancestry truncation (λ_GC_=1.02, FPR=0.050) or normalization (λ_GC_=1.00, FPR=0.051) in Figure 4C. On data simulated under a mix of the null and alternative, performance is similar for original (TPR=0.087, FPR=0.050), truncated (TPR=0.094, FPR=0.051), and normalized data (TPR=0.105, FPR=0.051). Thus, ancestry distribution transformations improve the performance of ancestry-variance tests, though these ancestry distributions do not cause substantial miscalibration.

### Variance effects in UK Biobank

Individuals from the British Isles have fine-scale population structure which is evident in a large sample^53^. To investigate whether ADGLM can detect fine-scale population variance structure, we applied ADGLM to ~113K British-ancestry, deeply-phenotyped individuals from UK Biobank (UKB, Supp. Materials). We tested binary, ordinal, and quantitative phenotypes (scaled to have mean 0 and variance 1). We included assessment center, genotype array, sex, age, and PCs1-10 as mean effects and tested for population variance structure 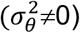 with ADGLM. We focus on genetic PCs1-5, which represent geographic population structure in UKB; PC1, specifically, is correlated with a geographic north-south cline^42^.

PC1 is associated (nominal *p*<0.05) with the phenotypic variance of 17 of 29 tested non-disease traits (Table S2), 12 of which are significant after Bonferroni correction (Table 2). Interestingly, 6 of these 12 associations are only with phenotypic variance, and not mean (absolute correlation of phenotype and PC1 < 0.01). Corpuscular hemoglobin has the strongest PC1 variance association among continuous traits (Figure S4). In addition, PCs2-5 are associated with the variance of 18 traits (Table S3), representing finer-scale population variance structure: PC3, which is also correlated with a north-south cline^42^, has the strongest of these variance associations.

**Table 2:** UK Biobank variance associations. Ancestry variance effect sizes estimates, standard errors, and p-values of associations of PC1 with phenotypic variance in UKB British-ancestry individuals. Associations are significant at a threshold of 0.05 after Bonferroni correction. Phenotypes followed by “(z)” are z-scores and starred phenotypes have an absolute correlated with PC1 that is greater than 0.01.

| Phenotype | Effect size ($\sigma_\theta^2$) | P-value |
| --- | --- | --- |
| Corpuscular hemoglobin | -0.025 ± 2.5E-03 | 7.75E-25 |
| Platelet volume * | 0.015 ± 2.5E-03 | 1.36E-09 |
| Platelet count | -0.012 ± 2.5E-03 | 7.13E-07 |
| Platelet distribution width * | 0.015 ± 2.5E-03 | 2.14E-09 |
| Erythrocyte (red blood cell) distribution width | -0.019 ± 2.5E-03 | 3.77E-14 |
| Leukocyte (white blood cell) count | -0.018 ± 2.5E-03 | 1.03E-13 |
| Educational attainment * | -0.020 ± 2.4E-03 | 5.73E-16 |
| Basal metabolic rate (z) | -0.024 ± 2.5E-03 | 3.41E-23 |
| Forced vital capacity (z) * | 0.012 ± 2.7E-03 | 8.89E-06 |
| Hair pigment | -0.023 ± 2.46E-03 | 1.28E-21 |
| Skin pigment * | -0.074 ± 2.45E-03 | 9.25E-199 |
| Skin tanning * | 0.068 ± 2.45E-03 | 4.02E-169 |

We also investigated whether age and sex are associated with phenotypic variance because age varies non-linearly with several phenotypes and different sexes represent different environments^13^. Of the 44 phenotypes tested, 33 have age-variance associations, and 17 have sex-variance associations (Table S2). Overall, population variance structure, age- and sex-variance associations are prevalent in a large sample of British-ancestry individuals.

### Variance effects in admixed populations

For the remainder of this work, we focus on three admixed populations from two asthma and allergy studies: Mexicans and Puerto Ricans from GALA II, and African-Americans from the Study of African Americans, Asthma, Genes, & Environments (SAGE II). We analyzed asthma, allergy-related diseases (eczema, hives, rhinitis, rash, and sinusitis), lung function (FEV1, FVC), change in lung function after the first (*δ*_1_) and second (*δ*_2_) albuterol dose, BMI, height, and skin pigmentation (baseline melanin, tanning). We adjusted phenotypic means for age, sex, and ancestry (African and European ancestry fraction for Mexicans and Puerto Ricans; African ancestry fraction for African-Americans).

Using ADGLM ancestry-variance tests 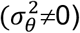, we find numerous associations (nominal *p*<0.05) of ancestry in Figure 5 (also see Tables S4-7), as well as age and sex (Tables S4-7) with phenotypic variance. The ancestry-variance effect sign for a given phenotype is the same across populations except for asthma, which has a negative African variance effect in Puerto Ricans and a positive African variance effect in Mexicans. To test for ancestry-variance heterogeneity, we performed a K-1 df LRT for a population with K ancestry fractions. Of the 14 phenotypes tested in three populations, 4 associations in 3 phenotypes are significant at a Bonferroni-corrected level of 0.05 (which is conservative because the phenotypes are highly correlated): asthma and *δ*_1_ in Puerto Ricans, and *δ*_1_ and *δ*_2_ in African-Americans. In addition, six of the phenotypes have previously-documented ancestry-mean associations which are also detected as mean effects with ADGLM (*β*_*θ*_≠0): FEV and FEV1 in Puerto Ricans^54^, asthma in Mexicans and Puerto Ricans^55^, baseline melanin^56,57^, *δ*_1_, and *δ*_2_ in African-Americans^58^.

**Figure 5:**
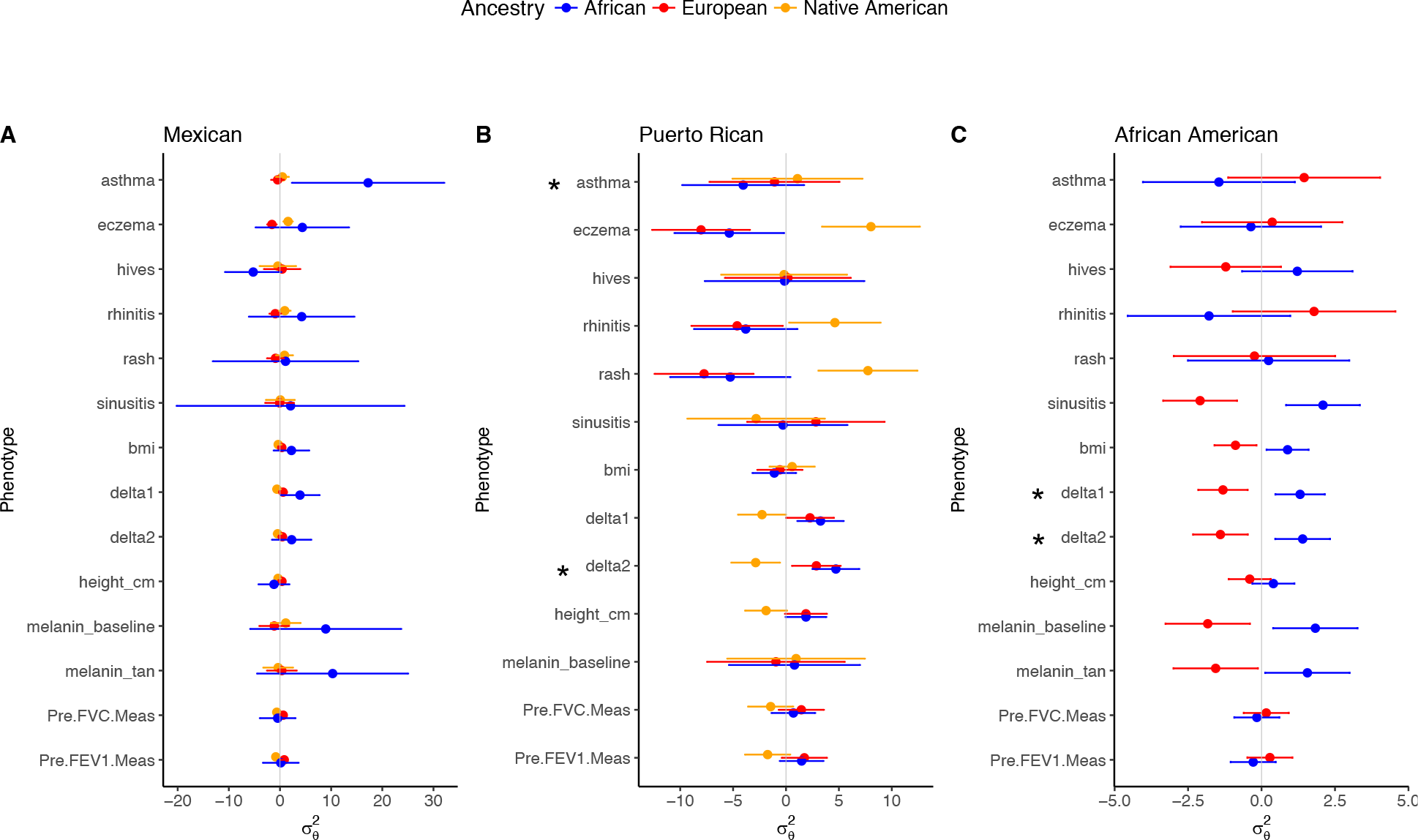
Ancestry-variance associations in admixed populations. Associations of African, European, and Native American ancestry with phenotypic variance in (A) Mexicans, (B) Puerto Ricans, and (C) African Americans. Points indicate estimates of ancestry variance effects, bands represent 95% confidence intervals, and starred phenotypes have significant ancestry-variance heterogeneity after Bonferroni correction (*p* < 0.05). The gray vertical line denotes 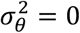.

### ADGLM GWAS in admixed populations

We tested for genetic associations (*β*_*g*_≠0) of common SNPs (MAF>0.05) in the admixed datasets above using both ADGLM, which corrects for population variance structure, and LR+PC, which does not. We represent ancestry using two ancestry fractions (African and European) for Puerto Ricans and Mexicans, and one (African) for African-Americans. Effect sizes for dichotomous traits (such as eczema) cannot be compared directly because they were obtained through probit regression. We discover two novel SNP associations with ADGLM, neither of which is significant with LR+PC: in Table S8, rs9808780 is associated with eczema in Mexicans and Puerto Ricans (*p*=1.53e-8) and rs113736578 is associated with rash in Puerto Ricans (*p*=2.14e-8). We next compared ADGLM to LR+PC at GWAS associations in the NHGRI catalog^48^, thinned to one SNP per locus. Since ADGLM is less inflated than LR+PC (Table S9), we genomic control^47^ adjusted test statistics when λ_GC_ >=1 to be maximally conservative. For the 46 GWAS SNPs in our datasets, 12 SNPs replicate with either test (*p*_*adj*_ < 0.05, Table 3;). Of these, 11 have a more significant p-value from ADGLM than LR+PC, indicating that ADGLM has better power to detect genetic associations than LR+PC.

**Table 3:** Replicated GWAS associations in admixed populations. Estimated main genetic effect sizes and p-values of replicated (*p*_*adj*_ < 0.05) NHGRI associations in admixed individuals (MX: Mexican, PR: Puerto Rican; FVC: forced vital capacity). ADGLM p-values are smaller than LR+PC p-values at 11 of 12 SNPs (starred).

| Population | Trait | SNP | LR+PC |  | ADGLM |  |
| --- | --- | --- | --- | --- | --- | --- |
| | | | Effect size ($\beta_g$) | P-value | Effect size ($\beta_g$) | P-value |
| MX | Eczema | rs17389644 * | 0.252 ± 1.51E-01 | 1.10E-01 | 4.90E-4 ± 2.15E-03 | 3.49E-02 |
| MX | Eczema | rs4796793 * | -0.412 ± 9.53E-02 | 3.85E-05 | -0.0074 ± 4.05E-02 | 2.76E-05 |
| MX | Melanin | rs16891982 * | 0.487 ± 1.70E-01 | 8.90E-03 | 0.591 ± 1.58E-01 | 8.47E-04 |
| MX | Tanning | rs12210050 * | 0.734 ± 3.32E-01 | 4.08E-02 | 0.748 ± 3.35E-01 | 2.50E-02 |
| MX | Tanning | rs7969151 * | 0.613 ± 2.70E-01 | 3.63E-02 | 0.803 ± 2.38E-01 | 4.10E-03 |
| MX | Tanning | rs154659 * | -0.393 ± 1.92E-01 | 5.73E-02 | -0.397 ± 1.78E-01 | 2.41E-02 |
| MX | FVC | rs9791644 | -0.148 ± 6.33E-02 | 2.65E-02 | -0.131 ± 6.32E-02 | 4.94E-02 |
| MX | Rhinitis | rs868688 * | -0.137 ± 6.22E-02 | 3.48E-02 | -0.002 ± 8.13E-03 | 2.25E-02 |
| PR | Eczema | rs13015714 * | -0.14 ± 7.29E-02 | 6.55E-02 | -3.22E-4 ± 7.52E-4 | 4.15E-03 |
| PR | Melanin | rs1834640 * | -0.374 ± 9.37E-02 | 1.81E-04 | -0.375 ± 9.32E-02 | 1.41E-04 |
| PR | Melanin | rs6142102 * | 0.244 ± 8.55E-02 | 7.02E-03 | 0.253 ± 8.40E-02 | 4.45E-03 |
| PR | Melanin | rs2675345 * | -0.514 ± 9.68E-02 | 8.84E-07 | -0.542 ± 9.48E-02 | 2.14E-07 |

### Variance QTL

We next tested for genetic variance associations 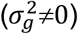 with ADGLM to find variance quantitative trait loci (vQLTs) in the admixed datasets above. For Mexicans and Puerto Ricans, we included genotype in the mean model and adjusted for ancestry, age, and sex in the mean and variance models; we did the same for African-Americans, adjusting for significant variance covariates (Table S6). We detect 17 vQTLs after genomic control adjustment (*p*_*adj*_ < 5e-8) in Table 4; the corresponding λ_GC_ values are in Table S10. The associations with hives in Mexicans, height in Puerto Ricans, asthma in African Americans, and tanning in African Americans are each detected in only one population. Of the 17 genetic variance associations, only 3 also have significant mean effects (rs1640275, rs117344403, rs55837614).

**Table 4:**
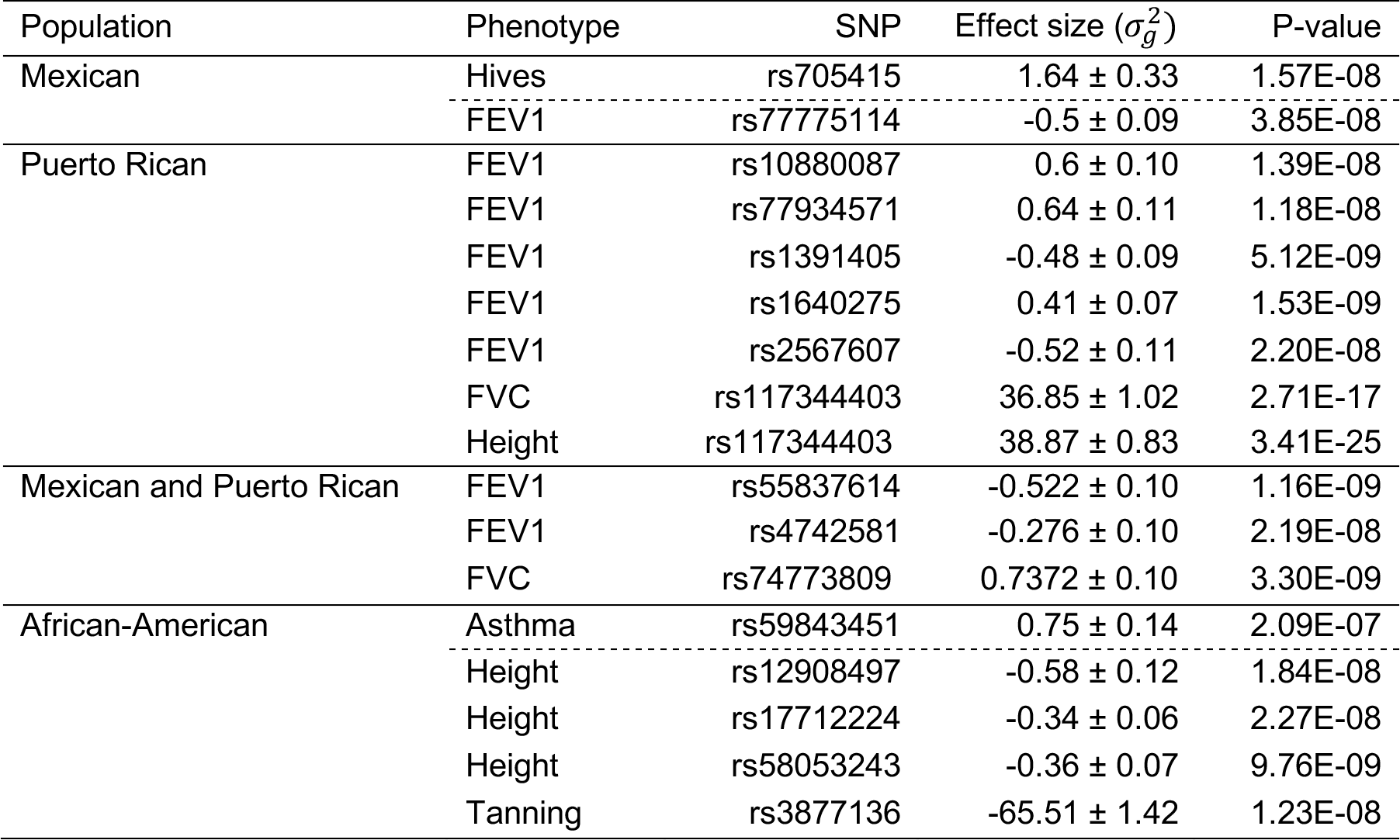
Variance quantitative trait loci in admixed individuals. Estimated genetic variance effects, standard errors, and p-values of vQTLs (*p*_*adj*_ < 5e-8, adjusted for genomic control) in admixed populations (FEV1: forced expiratory volume in 1 second; FVC: forced vital capacity). Tests of continuous phenotypes, which are below the dotted line within each population sub-table, were run on quantile-normalized phenotypes.

### Methylation association studies

DNA methylation, an epigenetic mark which is affected by environmental factors^27^, varies across disease phenotypes^28^ and ancestry^29^. To characterize the relationship of methylation and ancestry variance, we analyzed quantile-normalized methylation from 117 Mexican individuals. We adjusted for the mean effect of age, sex, ancestry, and asthma case status. We tested for ancestry-mean effects (*β*_*θ*_≠0) with ADGLM and LR+PC, as well as ancestry-variance effects 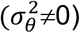 with ADGLM, resulting in Q-Q plots in Figure S5 and Manhattan plots in Figure S6. After Bonferroni correction, ADGLM identifies eight loci with ancestry-mean effects and 42 loci with ancestry-variance effects, 4 of which also have significant mean effects (Table S11). LR+PC, by contrast, only identifies one mean association, which is declared as a significant variance association, but not a mean association, by ADGLM.

## Discussion

In this study, we describe the presence of and discuss the importance of population variance structure, the difference of phenotypic variance by population. To model ancestry-variance relationships, we developed a novel statistical framework, the ancestry double generalized linear model (ADGLM). Unlike existing variance models, ADGLM accounts for continuous and discrete definitions of ancestry, arbitrary covariates, and binary or continuous phenotypes. We used ADGLM to discover many ancestry-variance associations in a British-ancestry and admixed human populations for a wide range of binary and continuous traits, including diseases and methylation, many of which have been subject to natural selection^42,56,57,59,60^.

When ancestry is related to phenotypic variance, genetic association tests with standard population structure corrections (e.g. linear regression with principal components adjustment or linear mixed models) are miscalibrated as a function minor allele frequency. This miscalibration has been observed for binary traits and can be attributed to the inability of standard LMM to model differences in disease prevalence^21^. We additionally observed this miscalibration for continuous traits and showed that it is a consequence of unmodeled population variance structure. Though not always apparent in a genome-wide Q-Q plot, this miscalibration can be readily detected by our diagnostic test which operates on summary statistics. ADGLM addresses these problems and association tests with ADGLM are both calibrated and well-powered for simulated and empirical human data.

The numerous variance associations we observed imply that previously conducted GWAS using LR+PC or LMM have residual population variance structure. The impact can be substantial as demonstrated by the inflation of height GWAS test statistics in LDscore analysis^61^. Two recent studies^62,63^ also found evidence for incomplete population structure correction in large cohort studies, including UK Biobank; based on our analysis, this may be due in part to unmodeled population variance structure. If phenotypes and principal components are available, population variance structure can be detected with ADGLM. Though the association of the square of a centered, scaled phenotype with principal components implies population variance structure may exist^64^, it is not a direct test.

In addition to acting as a statistical confounder, population variance structure has important biological implications, including in medical genetics^64^. Intuitively, differences in phenotypic distribution between populations imply that the fraction of individuals in the phenotypic tails differs between populations, and as such, longer tails may indicate a greater disease burden. As we previously showed, small differences in phenotypic variance between sexes can create large differences in disease liability^13^. Here, we estimate different asthma African ancestry-variance effects for Mexicans (17.2 ± 7.6) and Puerto Ricans (−4.05 ± 2.9). Mexicans and Puerto Ricans living in the U.S. differ dramatically in their asthma prevalence (8% vs. 22%), which has been referred to as the “Hispanic Paradox”^65^. These ancestry-variance associations might partially explain this difference.

In the 1940s, Waddington proposed that phenotypic variability is under genetic control, biological systems evolve to maintain homeostasis under a certain range of environmental or genetic perturbations, and that changing the environment to be outside the normal range will shift an optimum that has been shaped by stabilizing selection over many generations^66^. Under this model, increasing admixture proportion or shifting environmental conditions will lead to an increase in population variance, a process known as decanalization^15^. Although this phenomenon is well-documented in other species, it has seldom been described in humans, where it has been proposed as an explanation for the dramatic increase in non-communicable complex disease prevalence^67^. The ability of ADGLM to identify population variance structure as a function of admixture proportion or specific environmental context offers a new avenue to identify factors associated with variance heterogeneity, and thus potential drivers of decanalization.

ADGLM could be used as a screening tool for signals consistent with the presence of epistatic or genotype-by-environment interactions. ADGLM vQTL, which are controlled for variance population structure, may point to unmodeled interactions (either GxG or GxE)^68^ that can then be tested for interactions with other loci or specific environmental variables^49^. Additionally, the presence of population variance structure may affect admixture mapping efforts^69^ and could be corrected for with ADGLM. Finally, we could reduce the running time of the ADGLM GWAS roughly to ordinary GWAS by fitting the background variance components only once^70^, rather than once per SNP.

In conclusion, we find pervasive population variance structure in multiple human populations. As human studies increase in size and diversity, models that account for population variance structure, such as ADGLM, will be required for interpretable association testing. ADGLM has utility in studies of non-human model systems and natural populations, which have differences in phenotypic variability among groups and variance effects. By focusing primarily on the effect of genetic variation on phenotypic mean and ignoring its effect on variance, we have been missing an important axis contributing to phenotypic variation and disease emergence. Modeling phenotypic variance with ADGLM will enable discoveries along this axis.

## Acknowledgments

We thank Simon Forsberg for data analysis support and Joel Mefford for insightful discussions. This research was conducted using the UK Biobank Resource under Application #30397. N.Z., S.M, A.D, and D.P were funded from NIH grants U011U01HG009080, 5R01HG006399, and 5K25HL121295. J.F.A was funded by NIH grant 5R35GM124881. This work was supported in part by the Sandler Family Foundation, the American Asthma Foundation, the RWJF Amos Medical Faculty Development Program, Harry Wm. and Diana V. Hind Distinguished Professor in Pharmaceutical Sciences II, National Institutes of Health R01HL117004, R01HL128439, R01HL135156, 1X01HL134589, R01HL141992, National Institute of Health and Environmental Health Sciences R01ES015794, R21ES24844, the National Institute on Minority Health and Health Disparities P60MD006902 RL5GM118984, R01MD010443 and the Tobacco-Related Disease Research Program under Award Number 24RT-0025, 27IR-0030.

## Web Resources

The URLs for data presented herein are as follows: ADGLM, https://github.com/shailam/adglm pylmm, https://github.com/nickFurlotte/pylmm

